# Accelerated plastic aging in suspension (APAS): A simple, reproducible approach for the generation of model micro- and nanoplastics through simulated environmental forces

**DOI:** 10.1101/2025.04.09.647903

**Authors:** Patrick M. Lelliott, Matthew Lindley, Mark Vidallon, Alison Hobro, Amber Delmas Eliason, Santiago Moreno-Caceres, Subharthe Samandra, Thomas Harrison, Rebecca Harper, Sylvain Trépout, Stephanie Devineau, Brad Clarke, Nicholas I Smith, Xiaowei Wang, David P. Marciano, Alexander R. Pinto

## Abstract

The environmental fragmentation of plastic waste leads to the formation of micro- and nanoplastics (MNPs), which pose serious ecological and human health concerns. Despite increasing interest in their biological effects, many studies rely on artificial, uniform particles that fail to mimic the diverse physical and chemical characteristics of real-world MNPs. To address this limitation, we developed the Accelerated Plastic Aging in Suspension (APAS) system—a scalable, reproducible method that mimics natural aging processes by combining ultraviolet (UV) radiation, thermal stress, and mechanical shear to generate environmentally relevant MNPs from commonly used polymers.

We used APAS to fragment polyethylene terephthalate (PET), polyamide 6 (Nylon), and polyacrylonitrile (PAN), and observed time-dependent degradation, including the spontaneous formation of nanoplastics (<100 nm). Flow cytometry revealed substantial increases in particle number and reductions in average particle size over 12 weeks. Imaging flow cytometry confirmed consistent generation of heterogeneous, irregular particles across replicate batches. High-resolution imaging via AFM, TEM, and SEM confirmed the presence of nanoplastics with textured and irregular morphologies.

Chemical characterization showed APAS aging altered particle surface charge and induced polymer-specific changes in autofluorescence and Raman spectral profiles, consistent with oxidative surface modifications. Laser Direct Infrared (LDIR) imaging further confirmed structural and chemical changes in polymer spectra post-aging. Functionally, under physiologically relevant shear flow conditions, endothelial cells internalized APAS-generated PET MNPs at significantly higher levels than polystyrene (PS) beads of similar size. Uptake was enhanced particularly under oscillatory flow, highlighting the influence of particle physicochemical properties on cellular interactions.

Together, these findings demonstrate the ability of the APAS system to produce complex and realistic MNPs for use in environmental and toxicological studies. The system enables generation of nanoplastics and supports more accurate modelling of biological exposure scenarios compared to conventional synthetic particles.

## Introduction

Pervasive and rapidly increasing plastic pollution has led to the generation of micro- and nanoplastics (MNPs), tiny fragments resulting from the breakdown of larger plastic items. These particles are now recognized as a global environmental concern due to their ubiquity and potential impacts on ecosystems and human health^1,2^. While efforts to understand the effects of MNPs on organisms have grown, significant gaps remain in accurately replicating real-world MNP exposure for experimental research^3^.

Existing studies often rely on the use of commercially manufactured, uniformly sized and shaped polymer beads, which fail to capture the heterogeneity of naturally occurring MNPs. Real-world MNPs exhibit a wide range of sizes, shapes, and surface chemistries, and their interactions with biological systems are heavily influenced by these properties^4^. Moreover, the combination of particles of different sizes can profoundly affect uptake and toxicity mechanisms^5^. This makes extrapolating findings from studies using uniform particles to real-world scenarios problematic.

Harvesting naturally occurring MNPs from environmental samples presents its own challenges, including limited reproducibility, contamination, and insurmountable technical constraints, especially for smaller particles such as nanoplastics. An alternative is controlled generation of MNPs through current approaches such as cryogenic milling^6^ or precipitation^7^. While a valuable resource for many applications relating to MNP research, these approaches again fall short of generating particles that faithfully represent the properties of real-world MNPs. Cryogenic milling exposes plastics to extremely low temperatures which increase brittleness and milling effectiveness; however, these temperatures do not occur naturally and therefore the fragmentation pattern, surface texture and size distribution of resultant MNPs may differ from those formed in natural conditions. Alternatively, solubilisation of polymers and controlled precipitation through transfer into different mediums at different flow rates enables precise control over particle size and properties, but again, does not represent natural formation processes.

The primary environmental factor driving the formation of MNPs is exposure to ultraviolet (UV) radiation^8^. UV mediated photodegradation alters the properties of plastics, making them prone to fragmentation and altering their surface characteristics^9,10^. UV irradiance causes the breakdown of polymer bonds, changing polymer structure and reducing polymer chain length. Along with photo-oxidation this alters the texture and chemical properties, such as surface charge, of particles. Together, these changes in properties can influence biological effects of MNPs^11^. Additives, such as antioxidants and UV stabilizers, can slow this process and prevent rapid photodegradation of the polymer matrix until the additives are oxidized, degraded, or released in the environment.

To replicate this process, we developed the Accelerated Plastic Aging in Suspension (APAS) system as a simple, high-throughput, reproducible approach for the generation of environmentally relevant MNPs for toxicity studies. This approach leverages UV-B photodegradation, thermal effects, and mechanical shear to mimic and accelerate the natural fragmentation of plastics. The APAS system replicates key environmental forces in a controlled laboratory setting, allowing for reproducible production of MNPs with diverse and environmentally relevant characteristics.

In this study, we demonstrate the capabilities of the APAS system to produce MNPs from common plastic polymers, including polyethylene terephthalate (PET, polyester), polyamide 6 (Nylon 6), and polyacrylonitrile (PAN, used in acrylic textiles). We provide detailed characterization of the particles, focusing on their size, shape, and surface properties, and assess reproducibility of MNP generation. We highlight the system’s ability to generate nanoplastics and demonstrate differential cellular uptake of particles compared with polystyrene (PS) beads, highlighting potential differences in their biological impacts.

## Methods

### Reagents

Powdered plastic polymers of Nylon 6 (Nylon) (Goodfellow, UK), Poly(acrylonitrile-co-methyl acrylate) (PAN) (Goodfellow), polyethylene terephthalate (PET) (Nanochemazone, Canada) were used as raw starting material for weathering and MNP generation. Full details of plastics are provided in Table 1.

**Table 1.** Plastic source material.

| Polymer | Common and alternate names | CAS No. | Supplier | Product code | Product form |
| --- | --- | --- | --- | --- | --- |
| Polyamide 6 | Nylon 6, polycaprolactam | 25038-54-4 | Goodfellow, UK | AM30-PD-000155 | Powder |
| Poly(acrylonitrile-co-methyl acrylate) (PAN) | Acrylic textile | 25014-41-9 | Goodfellow, UK | AN31-PD-000110 | Powder |
| Polyethylene terephthalate (PET) | Polyester | 25038-59-9 | Nanochemazone, Canada | NCZ-AE-227 | Powder |

### Accelerated Plastic Aging in Suspension (APAS) system design

Plastic degradation was modelled on the American Society for Testing and Materials international standard (ASTM) D7238-20^12^ which combines UV, heat and condensation to simulate plastic degradation during end-use conditions. In our design we simulated the UV component of natural sunlight using UVB-313 fluorescent lamps (Sankyo-Denki, Japan) and irradiated plastics suspended in Milli-Q water under constant agitation to mimic plastic exposure to shear forces such as those in waterways, oceans, and during laundering. Quartz flasks were used to maximise UV transmission to plastics and glass-coated magnetic stir bars were used to avoid any contamination from plastic-coated stir-bars (Figure 1).

**Figure 1.**
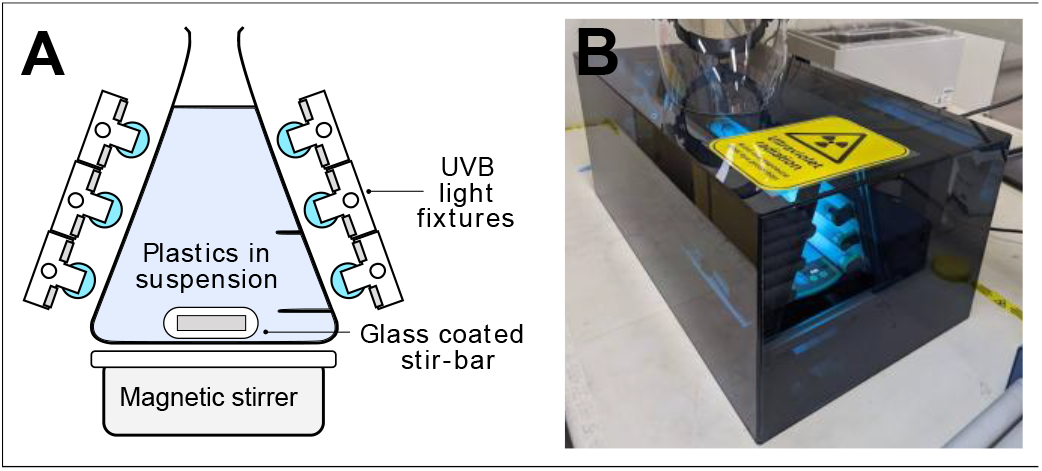
Accelerated Plastic Aging in Suspension (APAS) system design. Schematic (A) and photograph (B) of APAS system. Plastics in quartz flasks were suspended in Milli-Q water under constant agitation provided by a glass-coated magnetic stir bar rotated at 600rpm. Six UVB-313 lamps illuminated plastics at a maximal UV irradiance of 2250 µW/cm^2^ (accounting for 10% loss due to absorption by the quartz flask). A fan provided cooling, and the temperature of plastic suspensions was stable at 45-47 °C.

Plastic powders were washed in 80% v/v ethanol (Sigma-Aldrich, St Louis, MO). Ethanol was removed and plastics were resuspended in Milli-Q water at 100 mg/ml. For PET, 2% w/w bovine serum albumin (BSA, Sigma-Aldrich) was added to assist in particle dispersion. After 30 minutes of gentle, rolling agitation plastic dispersions were washed three times in Milli-Q water and resuspended at 10 mg/ml. A 4-position magnetic stirrer (LLG labware, Germany) was used to provide a mixing speed of 600 rpm for plastic suspensions. Three UVB-313 lamps were positioned either side of the flasks at a distance of 0-5 mm. Some variation in distance occurred due to differences in individual quartz flask dimensions. UV irradiance from 260 nm to 395 nm at this distance was 2500 µW/cm^2^, with 90% transmission through the quartz flask, as measured using a Center 532 UV light meter (Center, Taiwan). The apparatus was fully enclosed, with air circulation provided by an 80 CFM fan (Min quan, China) embedded at one end. The temperature of plastic suspensions stabilized at 45-47 °C during aging.

### Flow cytometry

Plastic suspensions were diluted to 1,000-10,000 particles per µL in Milli-Q water and filtered through a 70 µm mesh before measurement on an Aurora spectral flow cytometer (Cytek, Fremont, CA). Side scatter (SSC) gain for the violet laser (SSC-V) was set to 500 to maximise sensitivity, while SSC gain for the blue laser (SSC-B) was set to 2 to enable measurement of larger particles. A SSC-V of 500 was used as the threshold for event collection. PS beads 51 nm (Polysciences, Warrington, PA), 100 nm (Sigma-Aldrich), 180 nm, 510 nm, 1080 nm (Polysciences), 2.0 µm, 3.3 µm, 7.9 µm, and 16.8 µm (Spherotech, Lake Forest, IL) were used as size standards. Milli-Q water alone was used to determine background levels of background particles and noise, and this was subtracted from results. Data was analysed with Flowjo v10.9 (BD Biosciences, Franklin Lakes, NJ). To calculate particle size, a standard curve was prepared using 51 nm to 510 nm PS beads for SSC-V and 1.08 µm to 16.8 µm PS beads for SSC-B. Using a cut-off of 5M SSC-V, particles were separated into small and large populations, with sizes calculated using the SSC-V and SSC-B standard curves respectively. Overall mean particle size was calculated by combining the two means based on particle counts in each population.

### Imaging flow cytometry

Samples were filtered through a 40 µm mesh and analysed on an ImageStream^X^ Mark II instrument (Luminex, Austin, TX) using a ×60 objective. Events with a bright-field area >20 μm^2^ were collected. This cut-off was selected as shape characteristics of smaller particles could not be accurately measured. Imaging flow cytometry images were processed using IDEAS v6.2 (Luminex) and FlowJo v10.9 (BD Biosciences) as described previously^13^. Masking was optimised to identify the boundary of MNPs using the “Morphology” mask with a one pixel “erode” function applied.

### Electrophoretic Light Scattering (ELS)

To isolate the smallest fraction of plastic, suspensions were centrifuged three times at 20,000g for 10 minutes, each time the supernatant was collected, and pelleted plastics discarded. Plastics were then filtered through a 0.1 µm mesh and their surface zeta potential were analysed by ELS at 25 °C using a Malvern Zetasizer (Malvern Panalytical Ltd.).

### Light microscopy

Samples were mounted in Mowiol® 4-88 (Sigma-Aldrich) and imaged on an Olympus IX71 Microscope (Olympus, Japan) with a ×40 objective.

### Atomic Force Microscopy (AFM)

Morphology of plastic nanoparticles were evaluated using AFM based on a previously reported method^14^. Briefly, 5 µL plastic nanoparticle dispersions were drop casted onto clean glass slide and rapidly dried under a steam of nitrogen gas. Sample-coated slides were then imaged using a JPK Nanowizard 3 Atomic Force Microscope in intermittent contact (AC) mode with a Bruker NCHV cantilever (resonance frequency= 320 kHz; spring constant = 42 N m^−1^; tip radius = 8 nm).

### Scanning Electron Microscopy (SEM)

Plastic microparticle morphology was imaged using a JCM-7000 (JEOL, Japan) scanning electron microscope. For sample preparation, APAS plastics suspended in distilled water were dropped onto #1 cover glass (Matsunami) and allowed to air dry. Dried samples were then sputter coated with platinum to an estimated thickness of 4 nm using a JEC-3000FC auto finecoater (JEOL, Japan). Samples were then imaged with the SEM in high vacuum mode and at 15keV.

### Transmission Electron Microscopy (TEM)

TEM sample preparation and imaging protocol were adapted from previous studies^15,16^. Sample dispersions (3 µL) were pipetted onto a Parafilm®-covered glass plate. A continuous carbon film-coated, 300 mesh copper grid (EM Solutions) was incubated for 5 min on top of the sample droplet with the carbon face down. Following incubation, the excess sample was blotted away with filter paper and the grid was placed face down onto 10 µL droplets of 1% aqueous uranyl acetate. The grid was incubated on uranyl acetate for 30 s, after which the uranyl acetate was blotted away with Whatman™ filter paper, followed by drying prior to imaging. Stained sample-loaded grids were then imaged using a FEI Tecnai Spirit TEM at 120 kV.

### Raman microscopy and data analysis

Plastics suspended in distilled water were deposited onto quartz-bottom cell culture dishes (FPI, Japan) and allowed to air dry. Raman spectra of plastics before and after APAS were recorded using a Raman-11 (Nanophoton, Japan) microscope operating in line-imaging mode. Spectra were obtained using 532 nm excitation using a x 20 magnification objective (Nikon, Japan) and with a laser power of 2 mW at the sample plane. Each line in the Raman images was taken with 3 seconds exposure time, each image contained multiple particles, and triplicate images were recorded from different areas of the sample dish. The spectra were collected on a PIXUS 400 (Princeton Instruments, USA) camera via a spectrometer with 600 l/mm grating, resulting in a spectral range covering 524 to 2966 cm^−1^.

Raman images were concatenated to create final datasets of 210 × 385 pixels for each sample type. Pixels containing spectra originating from plastic were identified by thresholding the image based on scores from principal component 1 (PC1). To do this, principal component analysis (PCA) was performed. Spectra were processed using a 5 points Savitsky-Golay smoothing in the spectral domain and mean centring prior to PCA, and cross validation was performed using random subsets with 10 data splits and 5 iterations, and only those spectra that had a positive score on PC1 were retained for further analysis. These remaining pixels were then averaged together to produce one characteristic spectrum for each sample type. All processing steps and PCA were performed using the MIA and PLS toolboxes (Eigenvector Research Inc., USA) operating in MATLAB (Mathworks, USA).

### Laser Direct Infrared Spectroscopy (LDIR)

LDIR analysis was performed as previously described^17^. Polymers were placed on a MirrIR slide (Kevley Technologies, Parma, OH) and dried before being analysed on an 8700 LDIR Chemical Imaging System (Agilent Technologies, Santa Clara, CA). For each polymer four, 0.74 mm x 1.035 squares were analysed. The LDIR operates in the fingerprint region of polymers (1800 – 975 cm-1), where the greatest variation can be observed for most polymers^17^. To automatically classify microplastics, the particle analysis method was selected to analyse microplastics between <10 – 500 μm. only matches above 0.8 were classified as positive matches.

### PET labelling with iDye Poly Pink

APAS treated PET was filtered through a 40 µm and a 5 µm mesh before preheating to 70°C. iDye Poly Pink (Jacquard, Healdsburg, CA) was dissolved in 50% ethanol at 20 mg/ml. The iDye stock solution was added to the preheated PET sample for a final staining concentration of 2 mg/ml for 30 minutes at 70 °C, with agitation every 5 minutes. To remove unbound dye, ten volumes of 70 °C milliQ water were added and samples pelleted at 4,000g, repeated 3x. The median particle size was 0.48 µm as measured by flow cytometry.

### HLMVEC micro-nanoplastic uptake analysis by immunofluorescence microscopy & flow cytometry

Human Lung Microvascular Endothelial Cells (hLMVEC; Lonza, Switzerland, CC-2527) were cultured in VascuLife VEGF-Mv Endothelial media (LifeLine, Oceanside, CA, LL0005) to passage 7 and seeded on Ibidi µ-Slide 0.4 luer glass bottom slide (Ibidi, Germany, #80177) coated with collagen. Slides were incubated for 2 hours at 37 °C with 5% CO_2_ to allow cells to adhere. hLMVEC were preconditioned with laminar or oscillatory shear (5.0 dyne/cm^2^; 1 Hz) in parallel for 48 hours. APAS treated PET nanoplastics labelled with iDye Poly Pink (5 × 10^6^ pt/mL; 0.48 µm) and PS beads (5 × 10^6^ pt/mL; 0.5 µm Fluoresbrite YG Microspheres, Polysciences) were added and images were taken after 4 hours using a Keyence BZ-X800 fluorescence microscope (Keyence, Japan). After imaging, slides were washed to remove unbound particles and cells detached with TrypLE Express (ThermoFisher, Waltham, MA). Cells were fixed with 2% PFA and analyzed on a NovoCyte Advanteon VBR flow cytometer (Agilent Technologies) (flow rate 6 µL/min, 5.1 uM core diameter).

## Results

### Plastic fragmentation during Accelerated Plastic Aging in Suspension (APAS)

To produce MNPs exhibiting characteristics of MNPs found in real-world settings, we generated a system that exposes plastics to both UV radiation and shear stress to accelerate plastic aging and fragmentation (Figure 1; See Methods). This *Accelerated Plastic Aging in Suspension* (APAS) system effectively degraded powdered plastics into MNPs, as demonstrated with polyethylene terephthalate (PET), polyamide 6 (Nylon), and polyacrylonitrile (PAN). Representative images of plastic particles before and after aging confirmed significant degradation and fragmentation over 12 weeks of APAS (Figure 2A). PAN particles exhibited significant fragmentation and cracking after 2 weeks aging, and larger particles were nearly completely degraded by 4 weeks. PET and Nylon degraded more slowly, although an increase in smaller fragmented particles was evident after 4 weeks. All plastics seemed to form irregular films and aggregates with decreased contrast during aging, in particular, nylon seemed to completely degrade into irregular nanoplastics (< 1 µm) and films by 12 weeks, which were difficult to observe due to low contrast.

**Figure 2.**
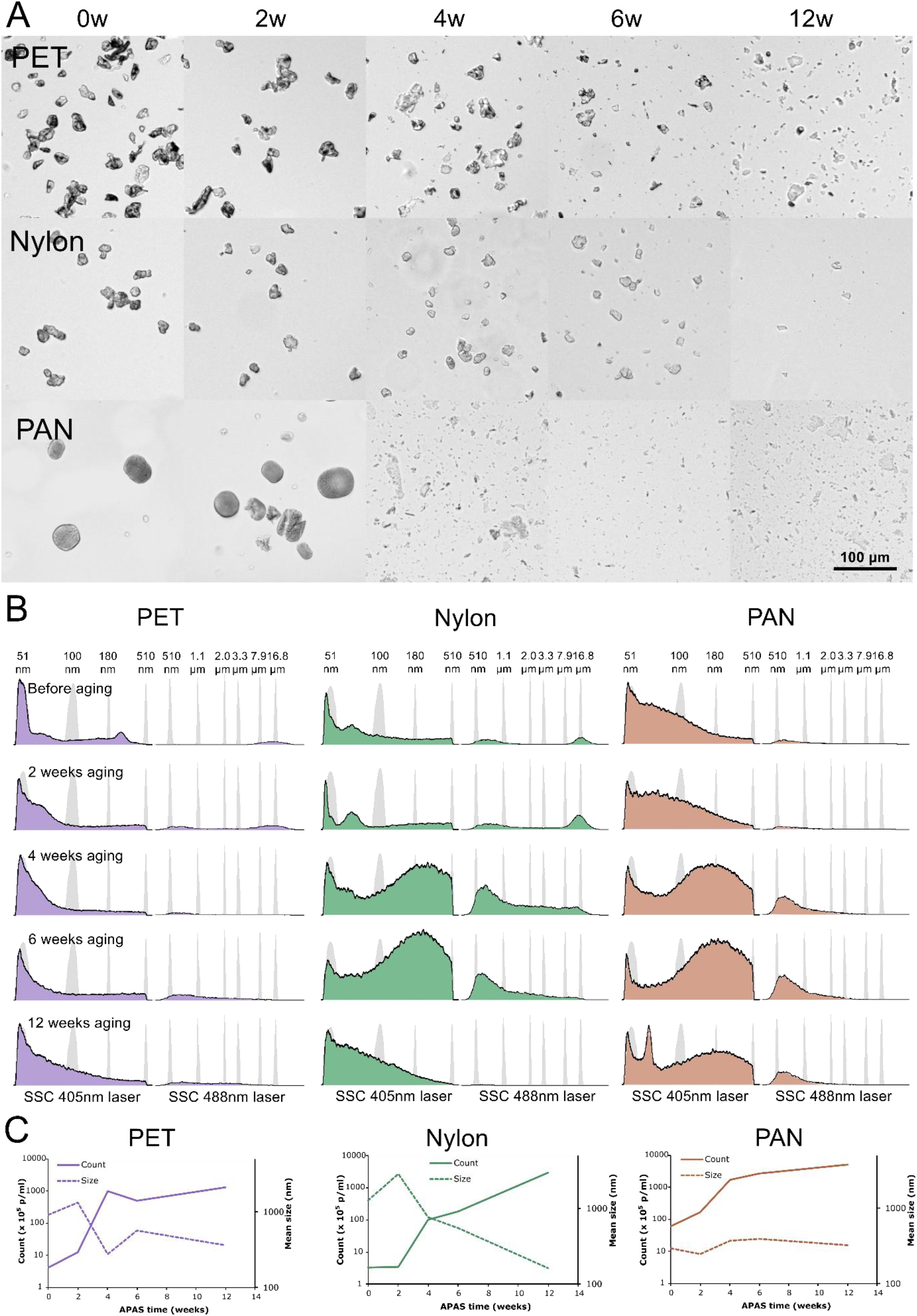
Breakdown of plastics under the Accelerated Plastic Aging in Suspension (APAS) system. (A) Representative images of polyethylene terephthalate (PET), polyamide 6 (Nylon), and polyacrylonitrile (PAN) powdered plastics before, and after 2-12 weeks of APAS. (B) Flow cytometry side scatter (SSC) histograms of plastics during APAS. Data is split, with smaller particles measured using 405nm SSC and larger particles with the 488nm SSC. PS bead size standards are shown in grey. (C) Particle count and mean size during APAS. Size was calculated using PS beads as reference.

To quantify particle count and size during APAS we developed a flow cytometry-based method to measure particles from ~50 nm to 70 µm. The lower limit was dictated by instrument sensitivity, while the upper limit was due to size constraints of the fluidics system. Our approach utilized side scatter (SSC) intensities produced by two separate lasers. To obtain the complete size range, one laser-detector pair was optimized to measure small particles with maximal sensitivity, while the other was adjusted to low sensitivity so that larger particles would not saturate the signal. Measurements of PS beads confirmed our ability to detect particles in this range and provided a calibration curve of particle size against SSC intensity which was used to calculate an estimation of MNP size (Figure 2B).

For all plastic samples, initial starting material contained significant amount of nanoplastics sized < 100nm. Although these particles contribute minimally to overall plastic mass, their small size and high number skew size distribution histograms. For PET, an initial peak close to 15 µm can be observed which decreases along with an increase in particles between 50-100 nm in size during aging. In contrast, both Nylon and PAN largely formed particles 200-500 nm in size, with nylon particles getting progressively smaller, reaching 100 nm and less as aging continues. Particle count increased rapidly during APAS, with a 2-3 order of magnitude increase over 12 weeks for all plastic types (Figure 2C). For PET and PAN particle number increased rapidly to 4 weeks then tapered off, while for nylon it continued to increase up to 12 weeks. Average particle size decreased during aging in PET and Nylon samples, while for PAN it remained stable. The later result is likely explained by the fact that larger PAN particles are unable to pass through the 70 µm sieve required for this analysis and therefore this size estimation does not capture their fragmentation during aging.

### APAS MNP morphology and reproducibility

Reproducibility is a critical component of approaches aimed at generating MNPs as a model for toxicological studies. To judge the consistency of MNP formation through APAS we analysed particle size and shape characteristics using imaging flow cytometry. We examined particles after 4 weeks of APAS, excluding those out of focus and with area less than 20 µm^2^, revealing heterogenous irregular shapes (Figure 3A). Importantly, comparison of four batches of MNPs revealed consistent trends in particle characteristics, including size (area, length and perimeter), shape (aspect ratio, shape ratio and circularity), and structure (contrast), supporting the reproducibility of the APAS system (Figure 3B).

**Figure 3.**
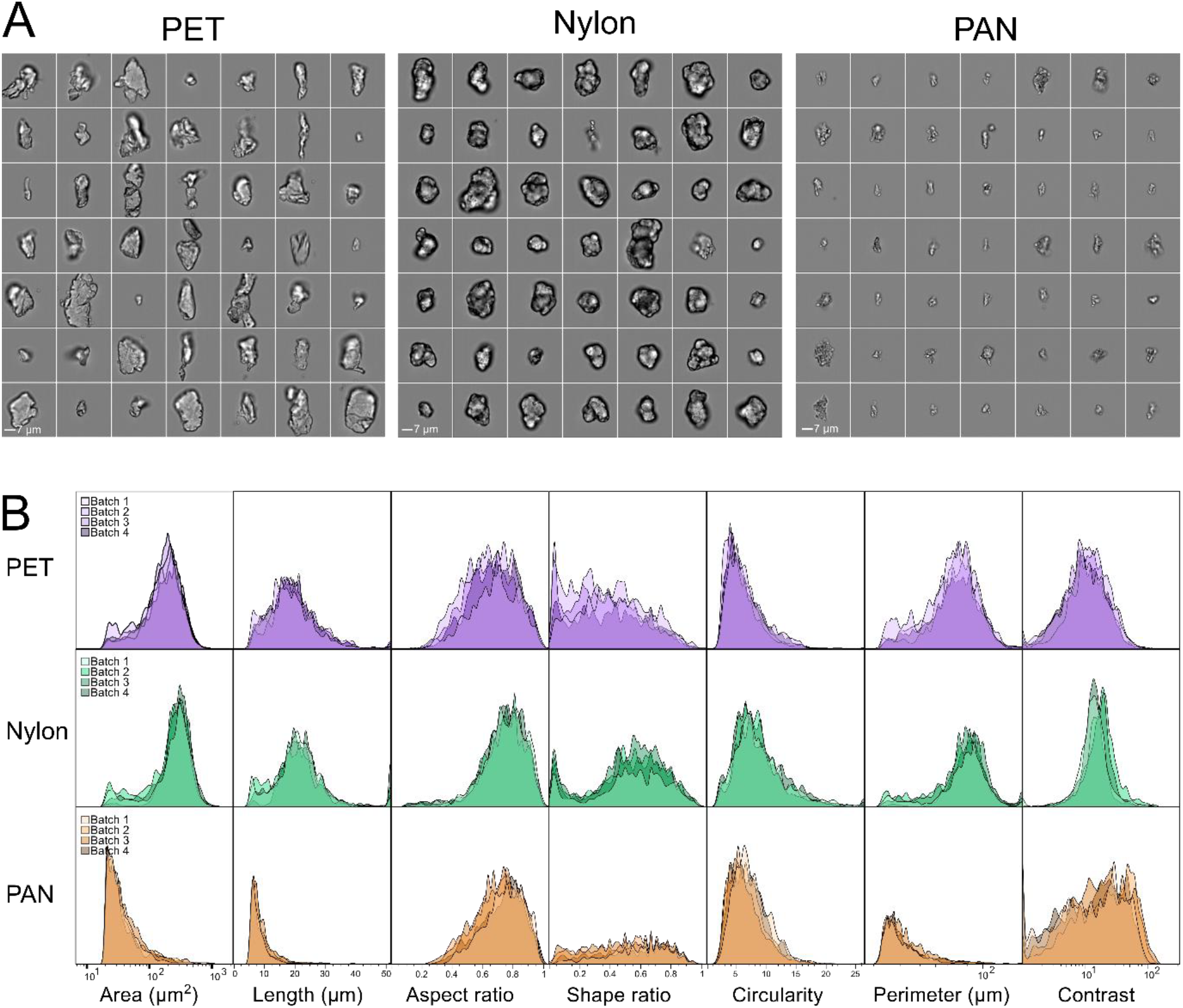
Batch comparison of particle size and shape produced using the Accelerated Plastic Aging in Suspension (APAS) system. (A) Representative imaging flow cytometry images of polyethylene terephthalate (PET), polyamide 6 (Nylon), and polyacrylonitrile (PAN) powdered plastics after APAS. Scale bar 7 µm. (B) Particle characteristics of four batches of microplastics subjected to APAS for 4 weeks masked and measured in the imaging flow cytometry analysis software IDEAS. Area = number of microns squared within the mask, length = longest part of the particle accounting for curvature, aspect ratio = minor/major axis, shape ratio = minimum thickness/length, circularity = degree of deviation from a circle, perimeter = boundary length in microns, contrast = image sharpness. A lower cut-off of 20 µm2 was used for all analysis.

### Generation of nanoplastics through APAS

The ability to produce and characterise nanoplastics is critical. We analysed the morphology of nanoplastics produced via APAS using multiple imaging modalities. Atomic force microscopy (AFM), transmission electron microscopy (TEM), and scanning electron microscopy (SEM) confirmed the presence of irregularly shaped particles <50 nm in size for all plastics studied after APAS (Figures 4A–C). Particle surfaces were highly textured and irregular.

**Figure 4.**
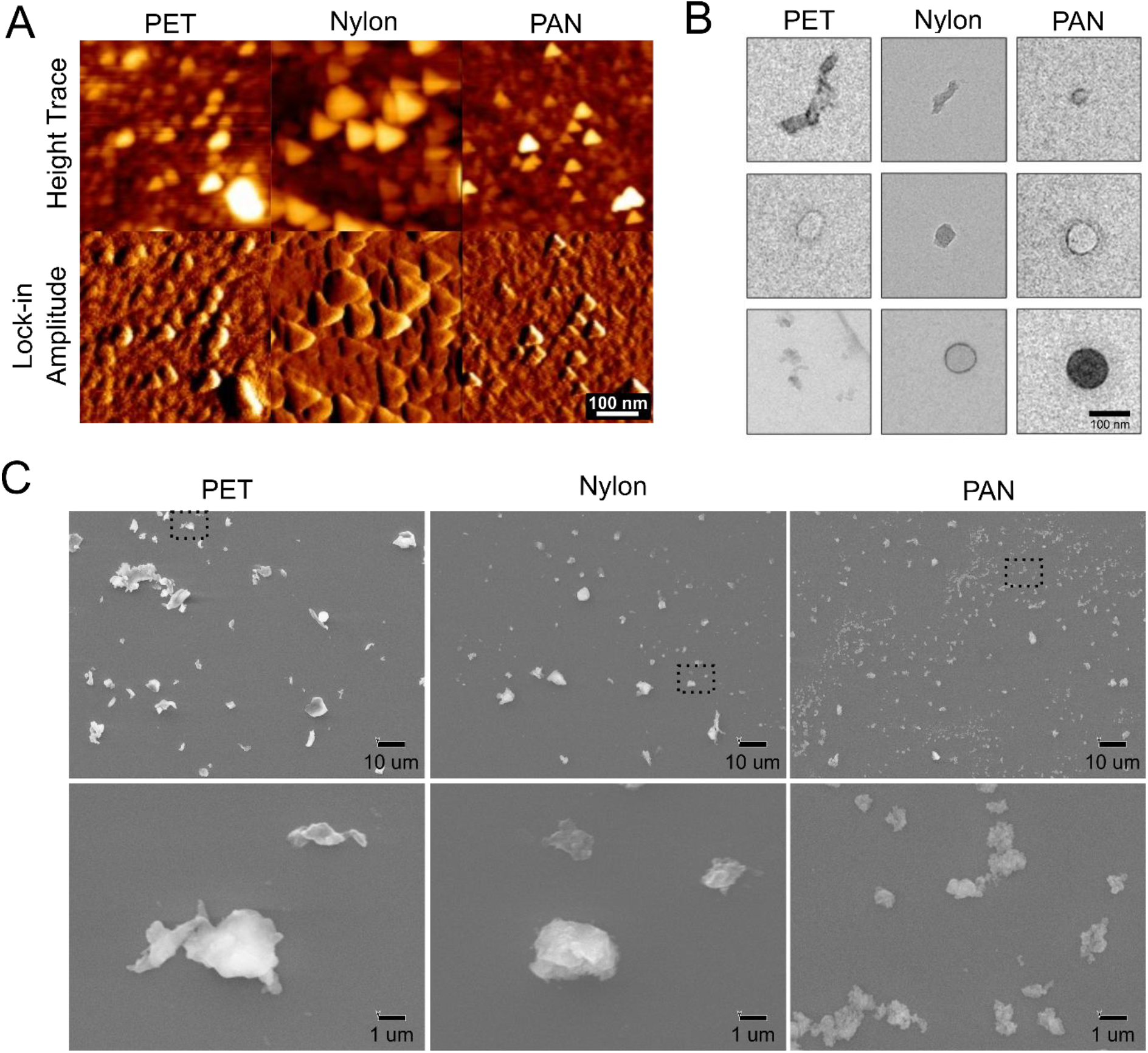
Nanoplastics produced through APAS. Representative AFM (A), TEM (B), and SEM (C) images of polyethylene terephthalate (PET), polyamide 6 (Nylon), and polyacrylonitrile (PAN) nanoplastics after APAS.

### APAS influences MNP chemical properties and autofluorescence

UV exposure causes oxidation and free-radical formation, therefore we next characterized chemical alterations caused by APAS. ELS measurement of zeta potential demonstrated that while all particles had near neutral charge, APAS increased surface charge over aging time for PET and nylon MNPs, consistent with altered particle surface chemistry (Figure 5A). Analysis of autofluorescence by imaging flow cytometry showed enhanced fluorescence intensity of aged PET and PAN plastics, with distinct spectral shifts depending on polymer type (Figure 5B). These changes reflect modifications in polymer surface chemistry induced by UV degradation during APAS. Consistent with these findings, Raman measurements of plastics before and after aging detected a substantial increase in baseline signal likely caused by fluorescence (Figure 5C) in the PET and PAN samples. The Raman spectral profiles obtained are all consistent with previously published Raman spectra of these plastics in fibre form^18^. Raman spectra from the nylon samples are quite similar before and after APAS, with little difference in the positions or relative intensities of the Raman bands. However, APAS does appear to affect the baseline of the Raman spectra. The spectrum of APAS nylon shows a slightly elevated baseline, which remains relatively uniform across the spectrum. The average Raman spectrum obtained from PAN before APAS clearly shows the intense band at 2239 cm^−1^, although there is a notable fluorescent contribution to the baseline of the spectrum that particularly affects the fingerprint region. After APAS, the average Raman spectrum of the PAN samples is dominated by a high intensity, dome-shaped baseline that eclipses the Raman bands in the both the fingerprint region and the high wavenumber regions, as well as leaving the 2239 cm^−1^ band barely visible, suggesting that the treatment leads to a large increase in fluorescence in the PAN samples. The effects of APAS are similar for the PET samples, where the spectrum obtained before treatment shows a sloped, although relatively straight, baseline with several clear bands visible in the fingerprint region. After APAS, this baseline has a dome-shape and the majority of the weak and medium intensity bands are no longer clearly distinguishable in the fingerprint region. Again, this increase in baseline intensity and shape change indicates an increase in the fluorescence contained within the PET samples after treatment.

**Figure 5.**
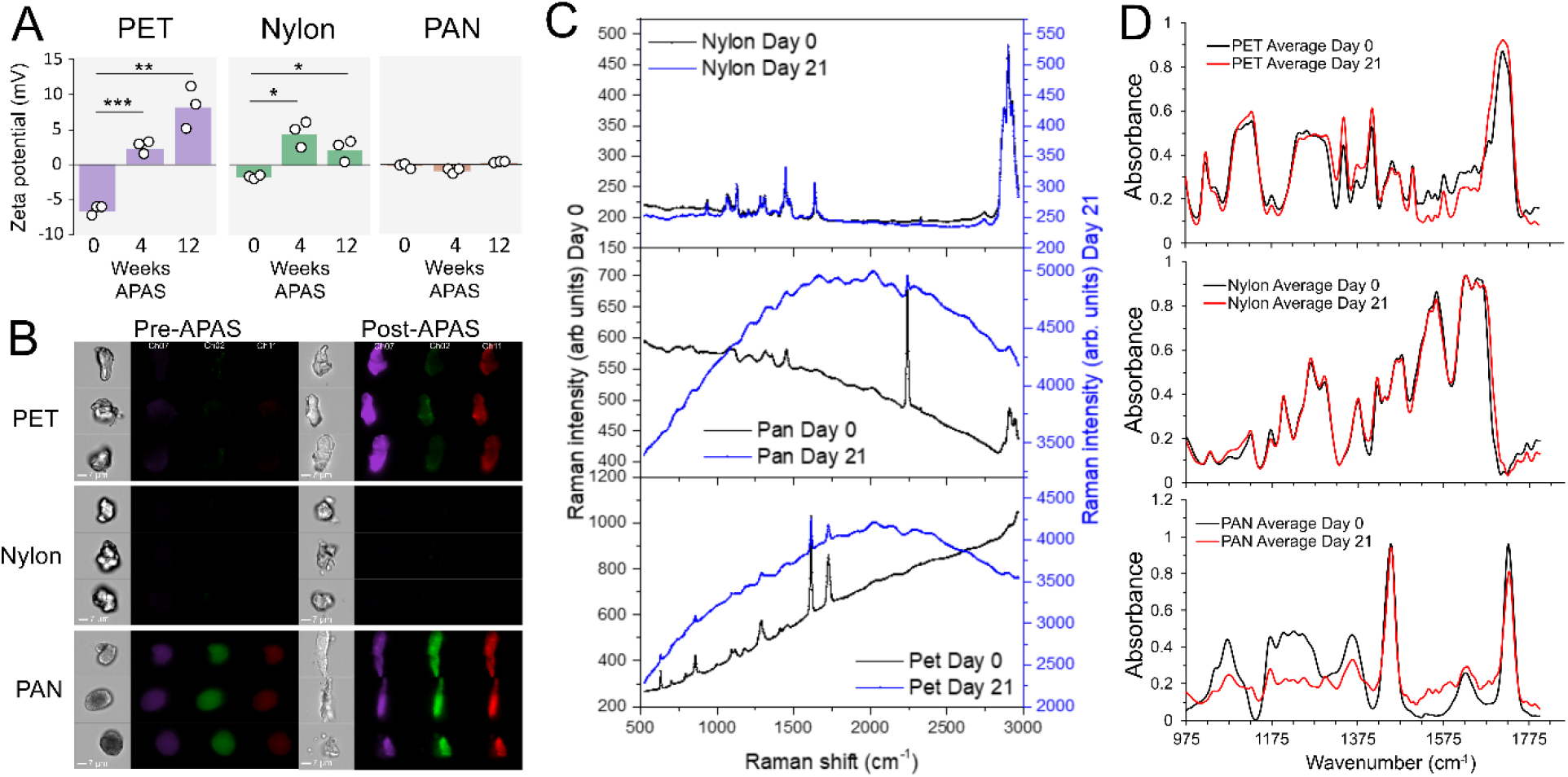
Influence of APAS on particle charge and spectral properties. (A) Zeta potential of polyethylene terephthalate (PET), polyamide 6 (Nylon), and polyacrylonitrile (PAN) powdered plastics before, and after 4 and12 weeks of APAS as measured by DLS. (B) Autofluorescence of plastics before and after 4 weeks of APAS measured by imaging flow cytometry. Ch07 excitation 405nm laser, emission filter 435-505nm. Ch02 excitation 488nm laser, emission filter 480-560nm. Ch11 excitation 405nm and 642nm laser, emission filter 642-745nm. (C) Average Raman spectra of PET, Nylon, and PAN particles at day 0 (black) or at day 21 (blue) of APAS. Raman intensity scales for day 0 samples are shown in black on the left-hand axis and Raman intensity scales for day 21 samples are shown in blue on the right-hand axis. Number of spectra included in the averages: Nylon day 0 n= 35344; Nylon day 21 n= 33488; PAN day 0 n=50072; PAN day 21 n=52551; PET day 0 n=34706; PET day 21 n=37403. (D) Average LDIR spectra of PET, Nylon, and PAN particles at day 0 (black) or at day 21 (red) of APAS.

We next examined plastics using Laser Direct Infrared (LDIR) imaging, a common approach used to identify and characterise MNPs. The LDIR system classified the majority of particles consistent to their polymer type both before and after APAS. However, there were notable changes in their IR spectra consistent with aging induced chemical changes (Figure 5D). For Nylon, although the spectra are largely aligned, slight variations are noticeable between 1575–1680 cm^−1^ and 1475–1575 cm^−1^. In the first region, the APAS nylon spectrum appears broader, suggesting a structural change caused by UV degradation. In the second region, a slight decrease in peak intensity is observed, indicating slight oxidation, which probably suppressed the peaks in this range^19^. For PET, APAS resulted in a significant increase in intensity between 1675 and 1775 cm^−1^, indicating an increase in C=O (carbonyl) groups, likely resulting from PET oxidation. In contrast, slight decreases in intensity were observed between 975–1175 cm^−1^ and 1475–1675 cm^−1 20^. The aromatic rings of PET typically vibrate in the 1500–1600 cm^−1^ region, and their deformation during the APAS process may have led to a reduction in vibrational intensity in this range^21^. For PAN, the peak associated with C=O is lower in intensity compared to that of the raw standard, which deviates from the trend observed in other polymers. In the region between 975–1380 cm^−1^, both peak shifting and broadening are evident, indicating substantial structural changes.

### Shear flow conditions differentially alter endothelial cell uptake of APAS MNPs and PS beads

We anticipated that MNPs generated by APAS would exhibit different biological interactions compared to PS beads. We particularly wanted to assess how particles would interact with vascular cells under varying physiological conditions. We therefore analysed particle uptake by endothelial cells under experimental blood flow conditions. Cell monolayers were co-exposed to PET APAS MNPs and PS beads of similar size and at equal particle concentrations under static (S), laminar shear stress (LSS) or oscillatory shear stress (OSS) conditions. Uptake was monitored by fluorescence microscopy directly (Figure 6A-B), or via cell detachment and flow cytometry (Figure 6C-D). LSS dramatically decreased uptake compared to S conditions, while for OSS, designed to mimic disturbed flow seen at vessel bifurcations, uptake of both beads and APAS MNPs increased. In all flow conditions PET APAS MNPs adhered to cells substantially more compared with PS beads (7 to 20-fold increase). Notably, APAS MNP uptake increased more than 9-fold under oscillatory flow, while bead uptake only increased 4-fold. Overall, suggesting increased interaction and uptake by endothelial cells, particularly in flow conditions, due to the chemical and physical properties of the APAS MNPs.

**Figure 6.**
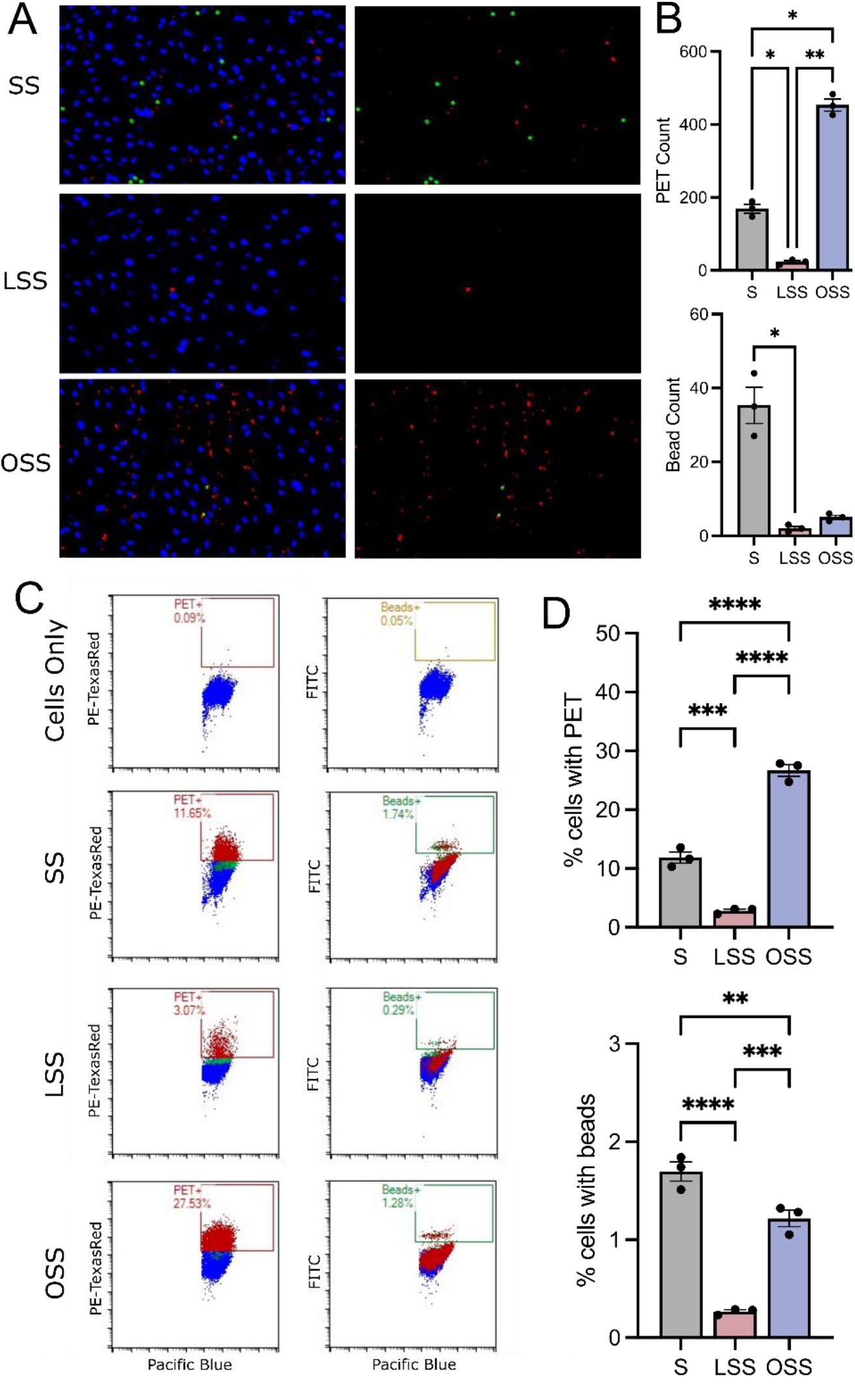
Uptake of MNPs by hLMVEC under different shear flow conditions. (A) Immunofluorescence of hLMVEC (DNA stain NucBlue, blue) after 4 hours of exposure to APAS PET MNPs (red) and fluorescent PS beads (green). (B) Quantification of microscopy results. (C) Representative flow cytometry plots of cells after wash and detachment identified based on NucBlue fluorescence on the Pacific Blue channel with PET and PS bead uptake measured by fluorescence on the PE-TexasRed and FITC channels respectively. (D) Quantification of flow cytometry results. S = static, LSS = laminar shear stress, OSS = oscillatory shear stress. * p < 0.05, ** p < 0.01, *** p < 0.001, **** < 0.0001.

## Discussion

The APAS system provides a novel and reproducible approach for generating environmentally relevant MNPs under controlled laboratory conditions. By combining UV photodegradation, hydrodynamic shear forces, and moderate thermal conditions (45 °C), the APAS system simulates key environmental processes that drive plastic degradation. This study highlights the system’s ability to produce MNPs with diverse but reproducible characteristics, bridging a critical gap in current methods for studying the biological and ecological impacts of plastic pollution.

The physiochemical properties of MNPs dictate their interactions with biological systems, such as cell uptake^4^, and will ultimately determine toxicity. Determining these characteristics in naturally occurring MNPs is a major technical challenge, particularly for smaller nanoplastics^22^. One strategy to estimate and reproduce the character of real-world MNPs is to examine the major forces driving plastic degradation in natural environments. Plastics are by design, highly resistant to degradation, however, they are susceptible to UV photodegradation, and this is widely considered the most significant factor driving degradation and consequent MNP formation in the environment. UV exposure causes photooxidation and polymer chain scission at the polymer surface^23^, in turn causing chain rearrangement and crosslinking, decreasing flexibility^24^. This can also lead to crack formation and propagation^25^, which splits particles into smaller sizes, a phenomenon we observed during PAN weathering. Importantly, UV not only causes degradation, but also influences the surface properties of particles generated, resulting in highly textured particles with high surface area and altered functional groups generated through generation of free radicals and oxidation^26,27^. These chemical changes influence surface charge and zeta potential^28^. In this study this was particularly evident for PET, in which APAS induced a shift from negative to positive zeta potential over time. Zeta-potential is well-known to influence cell-particle interactions, suggesting this is one way in which weathering may influence MNP toxicity. Notably, MNP weathering also influences surface interactions with environmental substances and their surface adsorption, so called “eco-corona” formation, which in-turn can alter zeta-potential^29^. Environmental plastics are often measured and characterized by their infra-red and Raman spectral properties. The rapid increase in autofluorescence observed during APAS aging is in agreement with measures from environmentally weathered samples^30^ and reenforces the importance of considering weathering effects when conducting MNP quantification.

While MNPs generated by the APAS system cannot fully replicate real-world MNPs, they are arguably a better representation compared to current alternatives, such as manufactured polymer beads, or MNPs produced through cryogenic milling^6^ or nanoprecipitation^7^. While these approaches are advantageous in that large amounts of material can be quickly purchased/generated with controlled properties, resultant particles lack the complex characteristics of those produced through UV mediated degradation. Cryogenic milling produces particles with angular shapes and unaltered chemical surfaces, while nanoprecipitation and polymer bead manufacture produces highly uniform spherical particles bearing little resemblance to environmental MNPs. Notably, although the particles generated by APAS were highly heterogenous, our imaging flow cytometry analysis suggests that batch-to-batch consistency is very high. For example, PET aging reproducibly produced particles of lower aspect ratio and circularity compared with nylon particles. Particles with lower aspect ratio (rod-like) shapes are better able to pass through intestinal barriers^31^, suggesting these differences in fractionation shape may influence particle biodistribution and toxicity.

Nanoplastics have been highlighted as of particular concern in terms of toxicity due to their higher surface area-to-volume ratio, enabling enhanced interactions with cells and biomolecules, and their small size which facilitates penetration into tissues and organelles. Depending on the route of degradation, the size distribution of nanoplastics and microplastics will change. We observed distinct degradation patterns for PET, PAN and nylon during APAS. PET directly formed nanoplastics <100 nm, while PAN and nylon initially formed larger nanoplastics ~200-500 nm. Attesting to the complexity of this fragmentation, nylon particles continued to breakdown, forming increasingly smaller particles, while for PAN this happened more slowly and a distinct population of nanoplastics sized ~75 nm formed. Under different conditions, such as cryogenic milling, the fragmentation pattern would be expected to differ, resulting in different biological interactions and toxicity due to size differences. Notably, the exponential increase in nanoplastics during degradation makes estimates of size distribution difficult, a single 20 µm spherical microplastic particle has enough mass to form 63 million 50 nm nanoplastics. Because of this, even small amounts of surface degradation will skew the size distribution towards smaller particles. This becomes more evident as sensitivity for detection of smaller particles increases, with the potential for exponentially more nanoplastics to be detected. In our approach, we were able to detect relatively abundant nanoplastics ~50 nm even before aging, and these occurred despite extensive washing of starting material. These were not present in the Milli-Q blank samples, which had minimal levels of particles (these were subtracted from the presented data). A recent study has demonstrated that nanoplastics are shed spontaneously from plastics^32^, consistent with these findings.

The differential uptake of APAS-generated PET nanoplastics by endothelial cells, compared to PS beads, highlights the importance of using environmentally relevant models to study biological interactions. Oscillatory flow causes endothelial cells to increase expression of adhesion receptors and can significantly affect their surface charge by disrupting the glycocalyx layer, a negatively charged protective coating on the cell surface^33,34^. Under static conditions, PS beads were able to adhere to endothelial cells, however, in oscillatory flow bead adherence was reduced, suggesting that PS interactions with the vasculature may be low in vivo. In contrast, adhesion and uptake of APAS MNPs under oscillatory flow conditions was increased more than 2-fold compared to static, suggesting that their aged surface properties may promote interactions vascular endothelium under physiological conditions.

The APAS system remains an imperfect model of natural MNP formation, and it lacks factors such as microbial activity, seawater salinity, and exposure to chemical contaminants—each of which can influence MNP formation and behaviour. Suspension in water also substantially influences MNP formation and therefore APAS MNPs likely differ from MNPs created by degradation of plastic waste in terrestrial environments^35^. In addition, naturally formed MNPs will shift in environmental distribution based on factors such as buoyancy and aerodynamics, which will ultimately determine human exposure levels^36,37^. For example, larger particles of dense MNPs may settle to the ocean floor, while smaller particles are more likely to enter the atmosphere. As a major source of MNP exposure occurs through ingestion of seafood and other animals, patterns of MNP accumulation in tissues of food sources and accompanying alterations due to digestion and bio-corona formation will ultimately have a substantial impact on the characteristics of particles humans are exposed to. Estimates of the characteristics of MNPs that humans are exposed to are improving as analytical techniques are further developed, however, analysis of smaller nanoplastics in complex samples will likely remain a challenge for some time.

In summary, here we describe a simple, reproducible, and scalable method for generating MNPs using forces simulating those found in the environment. By producing particles with realistic heterogeneity and surface properties, APAS enables more accurate investigations into the environmental and biological impacts of MNPs. Expanding the capabilities of APAS to include additional environmental variables or different types of source material, such as fibres, will further enhance its utility, enabling a more comprehensive understanding of the effects of MNPs in complex ecosystems and their impact on human health.

## Acknowledgements

The authors thank Professor Katsumasa Fujita and Mr. Itsuki Yamamoto of Osaka University for providing and assisting with the scanning electron microscope. The authors acknowledge Microscopy Australia (ROR: 042mm0k03) at the Ramaciotti Centre for CryoEM, Monash University, enabled by NCRIS.

